# Assembly, comparative analysis, and utilization of a single haplotype reference genome for soybean

**DOI:** 10.1101/2024.04.26.591401

**Authors:** Mary Jane C. Espina, John T. Lovell, Jerry Jenkins, Shengqiang Shu, Avinash Sreedasyam, Brandon D. Jordan, Jenell Webber, LoriBeth Boston, Tomáš Brůna, Jayson Talag, David Goodstein, Jane Grimwood, Gary Stacey, Steven B. Cannon, Aaron J. Lorenz, Jeremy Schmutz, Robert M. Stupar

**Affiliations:** Department of Agronomy and Plant Genetics, University of Minnesota, St. Paul, MN, 55108, USA; HudsonAlpha Institute for Biotechnology, Huntsville, AL, 35806, USA; Department of Energy Joint Genome Institute, Lawrence Berkeley National Laboratory, Berkeley, CA, 94720, USA; Corn Insects and Crop Genetics Research Unit, US Department of Agriculture – Agricultural Research Service, Ames, IA, 50011, USA; Arizona Genomics Institute, School of Plant Sciences, University of Arizona, Tucson, AZ, 85721, USA; Division of Plant Science & Technology, C.S. Bond Life Sciences Center, University of Missouri, Columbia, MO, 65211, USA

**Author notes:** Author for correspondence, Department of Agronomy and Plant Genetics, University of Minnesota, 1991 Upper Buford Circle, 411 Borlaug Hall, St. Paul, MN 55108-6026.

## Abstract

Cultivar ‘Williams 82’ has served as the reference genome for the soybean research community since 2008, but is known to have areas of genomic heterogeneity among different sub-lines. This work provides an updated assembly (version Wm82.a6) derived from a specific sub-line known as ‘Wm82-ISU-01’ (seeds available under USDA accession PI 704477). The genome was assembled using Pacific BioSciences HiFi reads and integrated into chromosomes using HiC. The 20 soybean chromosomes assembled into a genome of 1.01Gb, consisting of 36 contigs. The genome annotation identified 48,387 gene models, named in accordance with previous assembly versions Wm82.a2 and Wm82.a4. Comparisons of Wm82.a6 with other near-gapless assemblies of ‘Williams 82’ reveal large regions of genomic heterogeneity, including regions of differential introgression from the genotype ‘Kingwa’ within approximately 30 Mb and 25 Mb segments on chromosomes 03 and 07, respectively. Additionally, our analysis revealed a previously unknown large (∼20 Mb) heterogeneous region in the pericentromeric region of chromosome 12, where Wm82.a6 matches the ‘Williams’ haplotype while the other two near-gapless assemblies do not match the haplotype of either parent of ‘Williams 82’. In addition to the Wm82.a6 assembly, we also assembled the genome of soybean line ‘Fiskeby III’, a rich resource for abiotic stress resistance genes. A genome comparison of Wm82.a6 with ‘Fiskeby III’ revealed the nucleotide and structural polymorphisms between the two genomes within a QTL region for iron deficiency chlorosis resistance. The Wm82.a6 and ‘Fiskeby III’ genomes described here will enhance comparative and functional genomics capacities and applications in the soybean community.

## INTRODUCTION

Reference genomes are essential for crop improvement as they equip the research community with a tool to develop genomic resources to understand economically important traits, deploy technologies to introduce genetic variation, and aid in molecular breeding (Varshney et al., 2020). Due to the limitations of early-generation sequencing technologies, previous reference genome assemblies were incomplete, containing unoriented sequence scaffolds, especially in regions with highly repetitive elements that are recalcitrant to sequencing (Gladman et al., 2023). This often left large unresolvable gaps in the assembly, mostly in heterochromatic and repetitive regions, which hindered the understanding of functional genomics in these regions (Wang et al., 2023).

Recent advances in sequencing, such as single-molecule and long-read technology, have delivered higher contiguity reads. The utilization of long read technology, combined with Hi-C data for phasing, and advancement in assembly algorithms, enables the generation of telomere-to-telomere (T2T) gapless chromosome level assemblies (Zhou et al., 2022). In the last three years, researchers have assembled complete T2T genomes for several plant species, including arabidopsis (Wang et al., 2022), banana (Belser et al., 2021), barley (Navrátilová et al., 2022), maize (Chen et al., 2023), rice (Huang, 2023), soybean (Garg et al., 2023; Wang et al., 2023), and watermelon (Deng et al., 2022). These T2T genome assemblies have become the gold standard for the new era of genomics. From the perspective of plant genomics, the era of gapless genomes has been utilized for gene discovery through improved annotation of nucleotide and structural variants, identification of tandemly duplicated genes, thorough characterization of centromeric and subtelomeric regions, and decoding highly repetitive segments of the genome (Belser et al., 2021; Deng et al., 2022; Chen et al., 2023; Gladman et al., 2023).

Soybean (*Glycine max* L. Merr.), one of the most important crops in the world, is a self-pollinating species with 20 chromosome pairs. Since 2010, several genomes have been assembled, predominantly using cultivar ‘Williams 82’ (Wm82) as the main reference genome for the soybean research community. Including the present study, there are now six different Wm82 genome builds available (Schmutz et al., 2010; Song et al., 2016; Valliyodan et al., 2019; Garg et al., 2023; Wang et al., 2023). These are highly valuable resources for soybean breeding and functional genomics, facilitating trait mapping, designing molecular markers, and other aspects crucial for soybean improvement.

The Wm82 cultivar was developed through a backcrossing strategy between genotypes ‘Williams’ and ‘Kingwa’, aimed at introgressing a Phytophthora root rot resistance locus (*Rps1_k_*), with ‘Kingwa’ as the donor parent (Bernard and Cremeens, 1988). This breeding process included generations of single seed descent prior to bulk harvesting in the later generations. Importantly, the plants selected at the end of single seed descent still maintain some heterozygous loci. Subsequent bulk harvesting in the later stages of the breeding process allowed the heterozygous loci to segregate and differentially fix these loci among different sub-lineages within the bulked population. This type of intracultivar heterogeneity is common in soybean breeding, as cultivars are essentially maintained as collection of near-isogenic sub-lines (Mihelich et al., 2020). In Wm82, (Haun et al., 2011) provided a thorough characterization of intracultivar heterogeneity at a molecular level. They used the SoySNP50K platform (Song et al., 2013) to genotype the parental lines and different individuals of Wm82, revealing that different Wm82 individuals exhibit residual genetic/genomic heterogeneity within specific chromosomal segments. At the genomic level, this heterogeneity is traced back to variable introgressions of the donor parent ‘Kingwa’ among the different Wm82 individuals (Haun et al., 2011). Thus, they concluded that ‘Williams 82’ as a cultivar consists of a slightly heterogeneous collection of sub-lines. While the vast majority of the Wm82 genome appears to be homogeneous among sub-lines, early assemblies contained a mosaic of ‘Williams’ and ‘Kingwa’ haplotypes within some genomic regions (particularly on chromosome 03), presumably stitched together using sequencing reads from different Wm82 sub-lines (Haun et al., 2011). Even in the era of T2T assemblies, independent assemblies of Wm82 may be predicted to have pockets of variation amongst one another, depending on where the seed/DNA was sourced. Moreover, establishment of a Wm82 accession that matches a near-gapless assembly would be valuable to the research community, as no such resource currently exists, to our knowledge.

Here, we report an updated near gapless assembly of ‘Williams 82’, named as version 6 (Wm82.a6), derived from a sub-line known as Wm82-ISU-01 (Haun et al. 2011). This assembly matches a newly established seed stock deposited at the USDA Soybean Germplasm Collection (PI 704477). Additionally, we assess the genomic variation among three different near-gapless Wm82 assemblies, revealing large segments of heterogeneity presumably attributable to the sequencing of distinct Wm82 sub-lines. The analysis of comprehensive and near-gapless assemblies provides new and robust information about genetic heterogeneity among soybean sub-lines, adding perspective and considerations when using these genomic resources. Finally, we demonstrate utilization of Wm82.a6 to discover variants in economically important traits. To this end, we performed a genomic comparison of Wm82.a6 with a newly developed genome assembly of soybean line ‘Fiskeby III’, specifically investigating a quantitative trait locus (QTL) interval associated with increased resistance to iron deficiency chlorosis (IDC).

## RESULTS AND DISCUSSION

### A more complete Wm82 reference genome for soybean

Here, we focus on updating the reference genome of soybean cultivar ‘Williams 82’ (version Wm82.a6) using the sub-line Wm82 ISU-01 (Haun et al. 2011). The new assembly can be accessed at Phytozome v13 (https://phytozome-next.jgi.doe.gov/info/Gmax_Wm82_a6_v1). This genotype has been established as accession PI 704477 in the USDA Soybean Germplasm Collection. We assembled the genome with 47.07x coverage PacBio HiFi and 175X Omni-C sequencing and polished the resulting contigs with 57X Illumina 2x150 reads. This effort produced a 1.01Gb assembly of the 20 chromosomes with only 36 contigs (Table 1). Among six recent ‘Williams 82’ reference genome assemblies (Schmutz et al., 2010; Song et al., 2016; Valliyodan et al., 2019; Garg et al., 2023; Wang et al., 2023), Wm82.a6 has the longest contig N50 at 44.5 Mb (Table 1, Supplemental Figure 1).

**Table 1.**
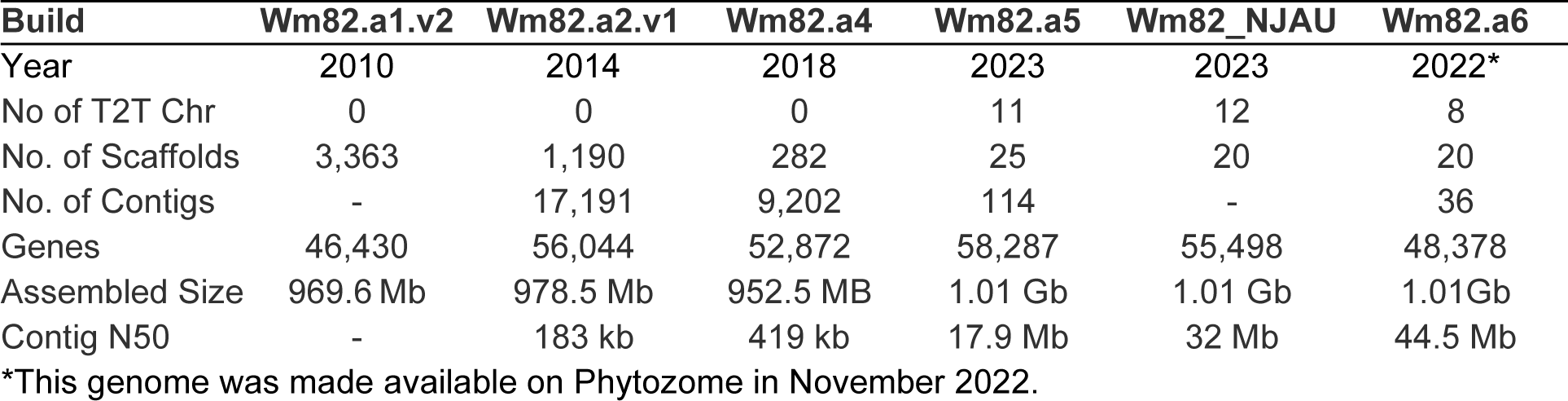
Genome assembly metrics of six versions of the Williams 82 reference genome build.

| Build | Wm82.a1.v2 | Wm82.a2.v1 | Wm82.a4 | Wm82.a5 | Wm82_NJAU | Wm82.a6 |
| --- | --- | --- | --- | --- | --- | --- |
| Year | 2010 | 2014 | 2018 | 2023 | 2023 | 2022* |
| No of T2T Chr | 0 | 0 | 0 | 11 | 12 | 8 |
| No. of Scaffolds | 3,363 | 1,190 | 282 | 25 | 20 | 20 |
| No. of Contigs | - | 17,191 | 9,202 | 114 | - | 36 |
| Genes | 46,430 | 56,044 | 52,872 | 58,287 | 55,498 | 48,378 |
| Assembled Size | 969.6 Mb | 978.5 Mb | 952.5 MB | 1.01 Gb | 1.01 Gb | 1.01Gb |
| Contig N50 | - | 183 kb | 419 kb | 17.9 Mb | 32 Mb | 44.5 Mb |

\*This genome was made available on Phytozome in November 2022.

Centromeric and telomeric repeats in the several Wm82 assemblies provide a way to assess assembly completeness. Centromeric arrays can extend several megabases in some species and some chromosomes, so can be challenging to accurately capture in genome assemblies. In the Wm82.a5, Wm82.a6, and Wm82.NJAU assemblies, both the centromeric and telomeric arrays are very close in size across all chromosomes (Supplemental Tables 1 and 2). The total count of 92-base CentGm-1 and 91-base CentGm-2 repeats is within 5% across all three assemblies -- the Wm82.NJAU assembly having 3% more repeats than the average across these assemblies, while Wm82.a5 and Wm82.a6 have 2% fewer. Consistent with Gill et. al. (2009), CentGm-1 was the dominant repeat element on the majority of chromosomes, with twelve chromosomes having predominantly CentGm-1 arrays and six chromosomes having predominantly CentGm-2 arrays. Considering telomeric completeness, all three assemblies have sizable arrays of the 7-base repeats - NJAU and Wm82.a6 respectively having 49,499 and 49,279 telomeric repeat units within 20 kb of chromosome ends, and Wm82.a5 having 43,178 such repeat units.

### Genes and repeat content of the newly assembled Wm82.a6 and synteny between six different Wm82 genome builds

We produced a well-supported and evidence-based gene model annotation to accompany our highly contiguous genome assembly. The 48,310 protein-coding genes in the Wm82.a6 gene annotation capture 99.5% of fabales_odb10 complete BUSCO genes (Benchmarking Universal Single-Copy Orthologs (Manni et al., 2021)). This is a marked improvement from Wm82.a5 (BUSCO = 96.6%), despite having nearly 10,000 fewer genes in the annotation (n. genes Wm82.a5: 58,105; Wm82.a6: 48,310, Supplemental Table 3).

The genomic landscape of Wm82.a6 is visualized in Figure 1, depicting genes (coding sequence (CDS) and introns), repeat content, and unannotated spaces. The gene and repeat content are shown in 900kb-overlapping 1Mb windows hierarchically in the following order, with the content abundance of each category in Mb: CDS (60.11), introns (135.76), Ty3-like retrotransposons (142.92), Copia-like retrotransposons (169.32), other repeats (146.4), centromere repeats Cen 91/92 (19.6), and unannotated sequence (337.02). Repeats including Ty3, Copia, and other repeats, constitute a significant fraction of the genomic landscape, while approximately one-third of the genome remains unannotated. As expected, the majority of genes are concentrated in the chromosome arms, while the repeat elements predominate in the pericentromeric region on the chromosomes.

**Figure 1.**
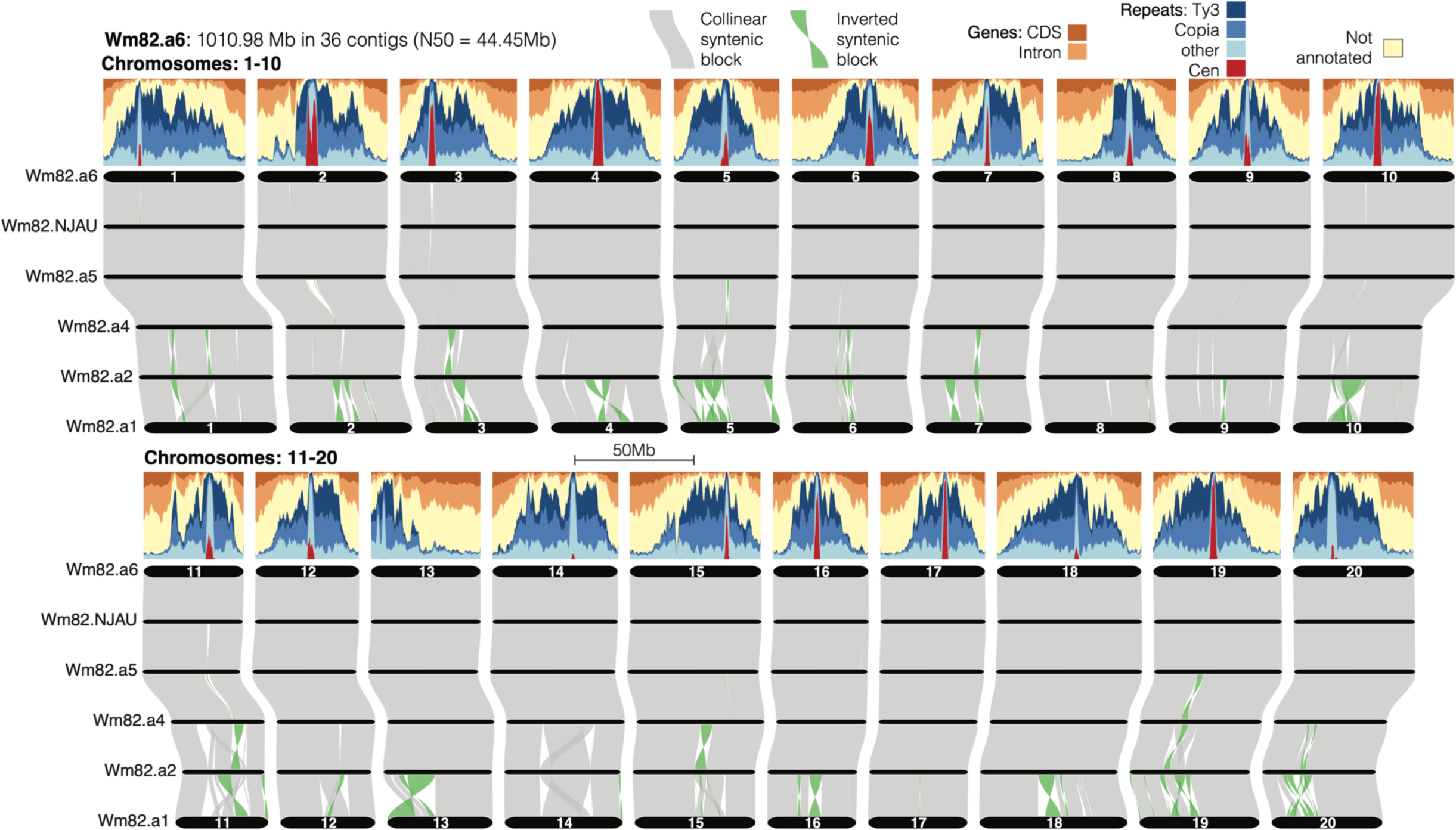
Genome landscape of Wm82.a6. Genic regions are concentrated on the chromosome ends, while repetitive regions are found in the pericentromeric regions. A comparison of six Wm82 reference genome builds reveals a high level of collinearity between the three newest assemblies (a5, NJAU, and a6) and in the genic regions on the older assemblies (a1-a4). Gray polygons represent collinear syntenic blocks, while green polygons represent inverted synthetic blocks.

Figure 1 illustrates the synteny between six different genome assembly builds, with Wm82.a6 as the reference. The results reveal a collinearity in the gene rich regions, which is consistent across all six genomes. The three newer near-gapless assemblies, Wm82.a5, Wm82.NJAU, and Wm82.a6, show a high level of collinearity except for a small region in chromosome 03. Notably, inversions are observed on the older genome versions (Wm82.a1 and Wm82.a2), but these have been resolved in the newer assemblies indicating an improvement in assembly quality over time. Furthermore, the near-gapless assemblies offer increased comprehensiveness and fill gaps in the genome, which may enhance structural variant detection and mapping efficiency, particularly in the heterochromatic regions.

Additionally, in the Wm82.a6 version, the gene name annotations remained consistent with the Wm82.a2 and Wm82.a4 assemblies, marking the first near gapless Wm82 reference genome to retain gene names from previous versions. This facilitates a smoother transition to the updated genome for the soybean research community, as gene names can be tracked across the different Wm82 genome assembly versions.

### Analysis of genomic heterogeneity among the six Williams 82 genome assembly builds

The soybean reference genome ‘Williams 82’ was previously found to exhibit genomic heterogeneity among individual stocks, as demonstrated by (Haun et al., 2011). A comparison of two Wm82 sub-lines (Wm82-ISU-01 and Wm82-SGC) found that the vast majority of chromosomes were homogenous, except for large blocks of heterogeneity observed on chromosomes 03 and 07 (Haun et al., 2011). While the SoySNP50K platform used in these analyses was able to demonstrate intracultivar heterogeneity, the full extent of heterogeneous regions among Wm82 sub-lines remained unknown. This is because these initial analyses were performed on a limited number of individuals and mapped onto an earlier version of the reference genome. Furthermore, the density of SNPs is sparse within the pericentromeric regions on the SoySNP50K panel.

Since 2010, different research groups have constructed updated versions of the Wm82 genome assembly (Schmutz et al., 2010; Song et al., 2016; Valliyodan et al., 2019; Garg et al., 2023; Wang et al., 2023). It is not always clear from where the seed sources used to generate these assemblies were derived. Moreover, it is possible that the type of heterogeneity described by Haun et al. (2011) may persist within specific seed stocks or collections, even the USDA Soybean Germplasm Collection, as described by Mihelich et al. (2020).

To test this assumption, we compared levels of heterogeneity among the six different Wm82 reference genome builds. To fully appreciate this question, it is important to note that ‘Williams 82’ was founded by four selected plants derived from a BC6F3 generation of ‘Williams’ (PI548631) × ‘Kingwa’ (PI548359) (Bernard and Cremeens, 1988). It is presumed that the major source of ‘Williams 82’ heterogeneity, particularly on chromosomes 03 and 07, is derived from differential recombination and segregation of ‘Kingwa’ haplotypes in Wm82 sub-lineages during the early (bulk-harvested) generations following the BC6F3 (Haun et al., 2011). Thus, in examining the heterogeneity among the six Wm82 genome assemblies, we were particularly interested in finding the genomic contributions of the respective parents, ‘Williams’ and ‘Kingwa’, to each assembly. We used the 50K SNP data of ‘Williams’ and ‘Kingwa’ to identify the SNP matches between the respective parental lines and each Wm82 reference genome across the 20 chromosomes (Figure 2a and Supplemental Figure 2). ‘Kingwa’ introgressions remain observable in all six reference genome builds, particularly on chromosome 03 where *Rps1k* is located, albeit to varying degrees, suggesting the presence of different haplotypes represented in the heterogeneous regions for each assembly.

**Figure 2.**
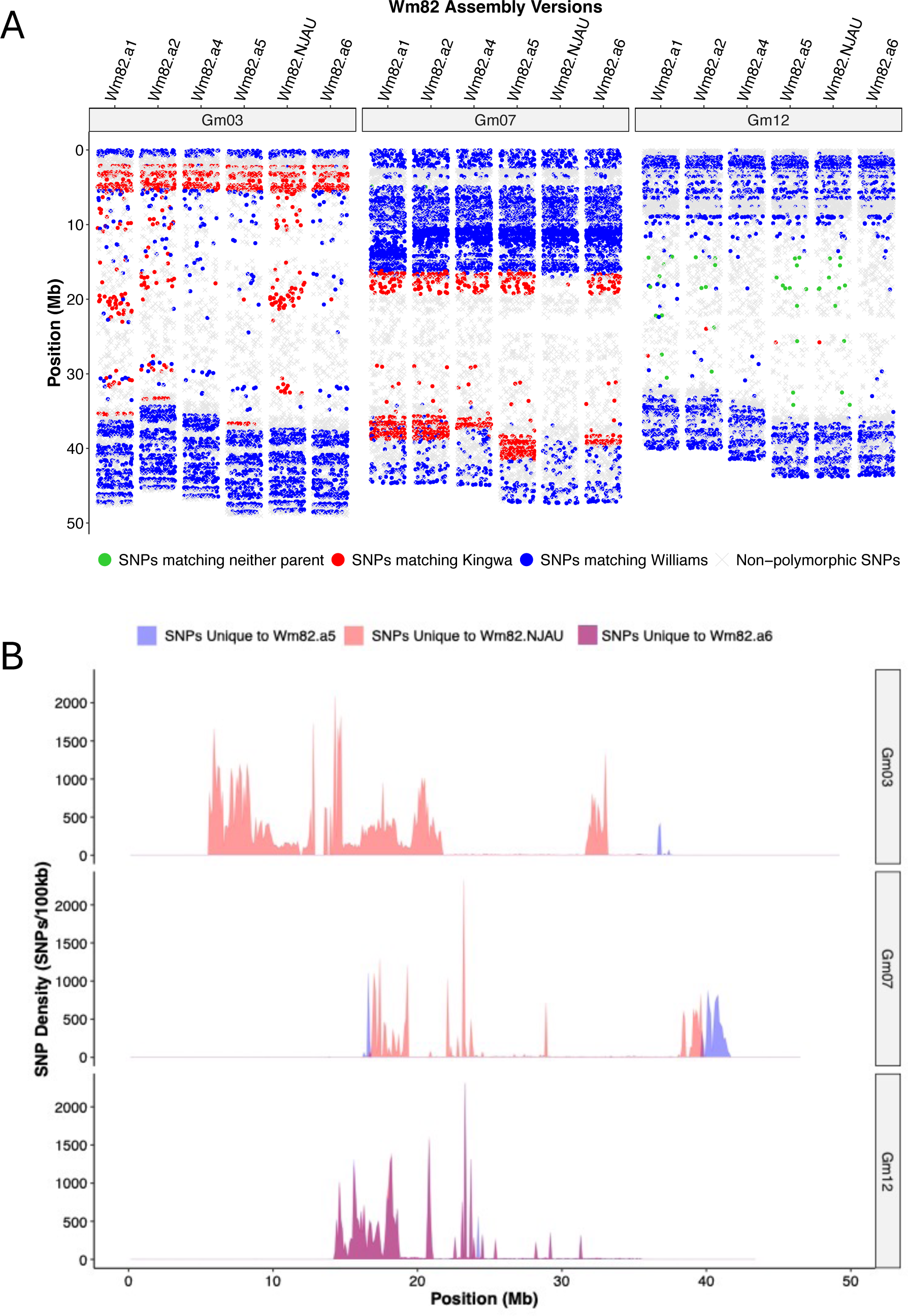
(A) SNP genotyping of Wm82 parental lines, Williams and Kingwa, compared to six different versions of Wm82 reference assembly showing heterogeneity and distinct haplotype at chromosomes 03, 07, and 12. (B) SNP density per 100 kb window of Wm82.a5, Wm82.NJAU, and Wm82.a6 on chromosomes 03, 07, and 12. SNP abundances unique to each genotype are shown according to the color code.

Recognizing that the 50K SNP data provides only a snapshot of the actual number of differing SNPs among the genomes, we further examined the extent of heterogeneity in these regions. We did comparative analysis of the three near-gapless Wm82 genome versions (Wm82.a5, Wm82.NJAU, and Wm82.a6), since these genomes are complete and contain more information in the pericentromeric regions, where the SoySNP50K array has a lower density of markers. Our findings revealed that the three assemblies clearly represent three different sub-lines of Wm82. We identified a total of 23,504 indels differing between Wm82.a5 and Wm82.a6, and 42,112 indels between Wm82.NJAU and Wm82.a6 (Supplemental Table 4). Altogether, we identified 97,030 SNPs unique to Wm82.NJAU, 32,034 SNPs unique to Wm82.a6, and 11,559 SNPs unique to Wm82.a5 (Table 2). The indels detected are distributed relatively evenly across 20 chromosomes for both genomes. However, the SNPs were enriched within specific regions, with notably higher numbers observed on chromosomes 03, 07, and 12, compared to the other chromosomes (Table 2). Small clusters of heterogeneity were also observed on chromosomes 14 and 17 (Supplemental Figure 3).

**Table 2.** Number of SNPs unique to each of the near-gapless assemblies summarized per chromosome.

| **Chromosomes** | **SNPs unique to Wm82.a5** | **SNPs unique to Wm82.NJAU** | **SNPs unique to Wm82.a6** | **SNPs unique for all three** |
| --- | --- | --- | --- | --- |
| Gm01 | 9 | 58 | 45 | 0 |
| Gm02 | 15 | 159 | 11 | 0 |
| Gm03 | 964 | 74,880 | 77 | 0 |
| Gm04 | 52 | 29 | 43 | 0 |
| Gm05 | 10 | 33 | 12 | 0 |
| Gm06 | 9 | 32 | 30 | 0 |
| Gm07 | 9,290 | 21,186 | 12 | 0 |
| Gm08 | 12 | 19 | 37 | 0 |
| Gm09 | 13 | 54 | 14 | 0 |
| Gm10 | 28 | 31 | 9 | 0 |
| Gm11 | 82 | 33 | 9 | 0 |
| Gm12 | 907 | 759 | 31,132 | 4 |
| Gm13 | 17 | 22 | 3 | 0 |
| Gm14 | 19 | 221 | 47 | 1 |
| Gm15 | 18 | 30 | 51 | 0 |
| Gm16 | 6 | 183 | 4 | 0 |
| Gm17 | 38 | 56 | 381 | 0 |
| Gm18 | 22 | 82 | 19 | 0 |
| Gm19 | 35 | 30 | 37 | 0 |
| Gm20 | 13 | 33 | 61 | 0 |
| Total | 11,559 | 97,930 | 32,034 | 5 |

### Chromosome 03

After closely examining the genetic heterogeneity among Wm82 sub-lines, it became apparent that the extent of ‘Kingwa’ introgression varies among the six reference genomes, particularly on chromosome 03. All genome versions show ‘Kingwa’ introgression in the upper portion of the chromosome (Figure 2A). This is expected, as this is the location of the *Rps1_k_* gene that was introgressed from ‘Kingwa’ during the breeding of Wm82, and is thus homogenous among all Wm82 individuals.

However, the six assemblies show significant disparities in their ‘Kingwa’ introgression through the lower and pericentromeric portions of chromosome 03 (Figures 2A and 2B). Wm82.a1, Wm82.a2, and Wm82.NJAU exhibit prominent tracts of ‘Kingwa’ introgression in the pericentromeric region. However, the Wm82.NJAU introgression does not extend as far as that of the Wm82.a1 and Wm82.a2 assemblies.

Among the three near-gapless genomes, there are 74,880 SNPs unique to Wm82.NJAU on chromosome 03, spanning positions ∼5.6-36 Mb (Figures 2A, 2B, and Table 2). The approximate size of introgression of ‘Kingwa” introgression in Wm82.NJAU is ∼30.4 Mb, with an average SNP density of 245 SNPs per 100 kb window. Wm82.a5 does not show a pericentromeric ‘Kingwa’ introgression, but does have a smaller segment introgression (∼3 Mb) in the lower portion of the chromosome, around position 35-38 Mb (Figure 2A and 2B). This indicates that different recombination events during the breeding of Wm82 (presumably following the single seed descent generations) produced a non-contiguous introgression on chromosome 03 in the sub-line used to develop the Wm82.a5 assembly.

Conversely, Wm82.a4 and Wm82.a6 show ‘Kingwa’ introgressions near the *Rps1k* region, but do not show evidence of introgression in the pericentromeric region nor the 35-38 Mb position. The relative similarity between Wm82.a4 and Wm82.a6 is not entirely coincidental, as genomic sequences from Wm82-ISU-01 were used to help assemble this region in Wm82.a4 (Valliyodan et al., 2019), and were the sole source of DNA for the Wm82.a6 assembly.

Notwithstanding the similarities of Wm82.a4 and Wm82.a6, comparisons between the six different Wm82 assemblies provide clear evidence that the different genome builds of Wm82 were derived from different heterogeneous sub-lines/individuals of Wm82.

### Chromosome 07

Similar to chromosome 03, the heterogeneous region on chromosome 07 exhibited different ‘Kingwa’ introgressions among the Wm82 assemblies. Wm82.a1, Wm82.a2, and Wm82.a5 showed large introgressions with similar boundaries. Wm82.a4 and Wm82.a6 showed slightly smaller ‘Kingwa’ introgressions, but still presumably spanned the pericentromeric region. A comparison of the near-gapless assemblies resulted in a total of 9,290 polymorphic SNPs unique to Wm82.a5, located at 39.7- 41.6 Mb, with a total introgression of ∼2 Mb.

In contrast to the other five assemblies, Wm82.NJAU showed almost no evidence of ‘Kingwa’ introgressions on chromosome 07 (i.e., almost all 50k SNP positions were non-polymorphic or matched the ‘Williams’ parent). Among the three near-gapless genomes, there are 21,198 SNP variants unique to Wm82.NJAU on this chromosome, primarily located at the positions ∼16.7-39.8 Mb, spanning an ∼23.1 Mb region (Figure 2B and Table 2). Similar to the heterogeneity observed in chromosome 03, it appears that chromosome 07 has at least three Wm82 haplotypes: one represented by Wm82.a5, one represented by Wm82.a6, and another represented by Wm82.NJAU.

### Chromosome 12

Prominent genomic heterogeneity was discovered among the Wm82 genome assemblies on chromosome 12, which was previously unreported. Upon comparing the three new assemblies, the Wm82.a5 and Wm82.NJAU share a similar haplotype containing approximately 32,000 SNPs around the pericentromeric region, specifically around positions 14.3 - 35.4 Mb, that are polymorphic with Wm82.a6 (Figure 2B and Table 2). This region spans approximately 20 Mb of heterogeneous region with an average SNP density of 150 SNPs per 100 kb. Upon closer examination, it appears that the Wm82.a6 SoySNP50K profile matches that of ‘Williams’ and ‘Kingwa’, which are not polymorphic in this region. Meanwhile, the two other assemblies, Wm82.a5 and Wm82.NJAU, have a subset of SoySNP50K markers in this region that do not match either of the Wm82 parental SNP profiles. At this stage, the origin of this “third party” haplotype is not clear. The simplest explanation is that the ‘Williams’ or ‘Kingwa’ plants used in the breeding of ‘Williams 82’ may be slightly heterogeneous to the individuals used in the SoySNP50K genotyping study (Song et al., 2015).

### The prevalence of genomic heterogeneity and limits of resolution

Perhaps the most important takeaway from the comparison of the near-gapless genome assemblies of Wm82 is that the greatest source of variation between the three genomes is not due to sequencing chemistry, assembly algorithms, or technical aspects. Instead, these three genomes are genomically distinct plants with biological differences due to differential genetic recombination and segregation. They are all ‘Williams 82’, but they are slightly different versions of ‘Williams 82’. The fact that they are different (while sharing the same name) is a byproduct of the method with which they were bred and the circumstance of having three different research groups perform their respective assemblies in parallel, but using different seed stocks/sub-lines to generate the DNA.

The main limitation of the current analysis is sample size. Three near-gapless genomes is a manageable number to analyze and present in an accessible format. However, had more near-gapless assemblies from more individuals of Wm82 been compared, it is reasonable to expect that other heterogeneous regions (perhaps even large heterogeneous regions) would have been identified in this study. Given the increased affordability and accessibility of near-gapless genome assemblies, such an analysis may be possible in the near future.

Looking beyond Wm82, approximately 4% of the USDA soybean germplasm collection exhibits inherent within-accession heterogeneity (Mihelich et al., 2020). Notably, the 4% estimate is based on a comparison of ∼3 individual plants per accession. A deeper sampling would likely increase this discovery rate. Furthermore, for a given accession, a comparison of the USDA stock with stocks held at other sites (e.g., breeding programs and/or individual research labs) could potentially reveal previously undiscovered genetic/genomic heterogeneity. Moreover, there are published reports of breeding programs selecting from within standing elite cultivars (Fasoula and Boerma, 2007; Sebastian et al., 2010), suggesting that intra-cultivar carries meaningful phenotypic variation, presumably caused by genomic heterogeneity.

To this end, we feel it is important that research groups develop their genomic resources (e.g., especially genome assemblies) from single individual plants and then maintain the seed stocks derived from those plants. This will allow other researchers to access the genetic resource (e.g., the seed stock) that matches the genomic resource (e.g, the genome assembly) for further research endeavors. As such, the Wm82.a6 genome is derived from a single plant (Wm82-ISU-01) and has a seed source deposited in the USDA soybean germplasm collection, under accession number PI 704477. Thus, the available biological resource matches the reference genome *per se*, providing a valuable resource for functional genomic studies.

### Genome assembly of ‘Fiskeby III’ and fine-mapping of an IDC QTL

In addition to the Wm82.a6 assembly, the ‘Fiskeby III’ genome was also assembled. This cultivar harbors various abiotic stress tolerance traits, including canopy wilt (Butenhoff, 2015), ozone (Burton et al., 2016), salt (Do et al., 2018), and IDC (Merry et al., 2019). ‘Fiskeby III’ was assembled from 133.95x PacBio reads, and the SNPs and indels were corrected using 50x Illumina reads (2x150, 400bp insert). Supplemental Table 5 and Supplemental Figure 4 shows the assembly statistics for the Fiskeby III genome. The Fiskeby III genome assembly compares favorably to earlier versions of the Wm82 reference genome (a.1-a.4), in terms of contigs, scaffolds, and genome size. Similar to Wm82.a6, the genic region (CDS and introns) for Fiskeby III also comprises less than 20% of the genome. Comparing genes predicted in Fiskeby III and Wm82.a6, there are significant differences in genes in the “defense response” GO category - but not unidirectionally. Defense genes have been lost and gained in both genotypes, as might be expected in this large and variable category of genes. Among 5,664 Fiskeby III genes without synteny-based orthologs in Wm82.a6, the genes are enriched in defense response (p-value 2.97 e-14; 86 genes). Among the 4,677 Wm82.a6 genes without synteny-based orthologs in Fiskeby III, the genes are also enriched in defense response (p-value 1.29 e-7; 70 genes).

The ‘Fiskeby III’ genome assembly can be employed for fine-mapping and characterizing the candidate genes of the abiotic stress resistance loci. The biparental population Mandarin (Ottawa) x Fiskeby III has previously been used to map an IDC QTL on chromosome 05 (Butenhoff, 2015). It was then narrowed down to a smaller interval spanning 137 kb and containing 17 gene models (Merry et al., 2019; Merry, 2020). To further fine-map the 137-kb interval, we selected heterogeneous inbred families (HIFs) to develop near isogenic lines (NILs) following (Tuinstra et al., 1997). We used the ‘Fiskeby III’ genome assembly to identify nucleotide variants across the region mapped for IDC resistance (Merry et al., 2019), and then designed KASP markers based on selected SNPs. These markers were used to screen for new recombinants. Seven families with recombination around the marker Gm05_62kb and Gm05_102kb were identified. In each of the seven NIL families, one line had the Mandarin (Ottawa) allele spanning all four markers (Mandarin_0-137kb), while the other line had the Fiskeby III allele on the left of a recombination region and the Mandarin (Ottawa) allele on the right (FiskebyIII_0-62kb) of the recombination region (see Figure 3A for a visual representation of the NIL pairs). After generating the NILs, these materials were harvested and planted in the IDC nursery across five different environments.

**Figure 3.**
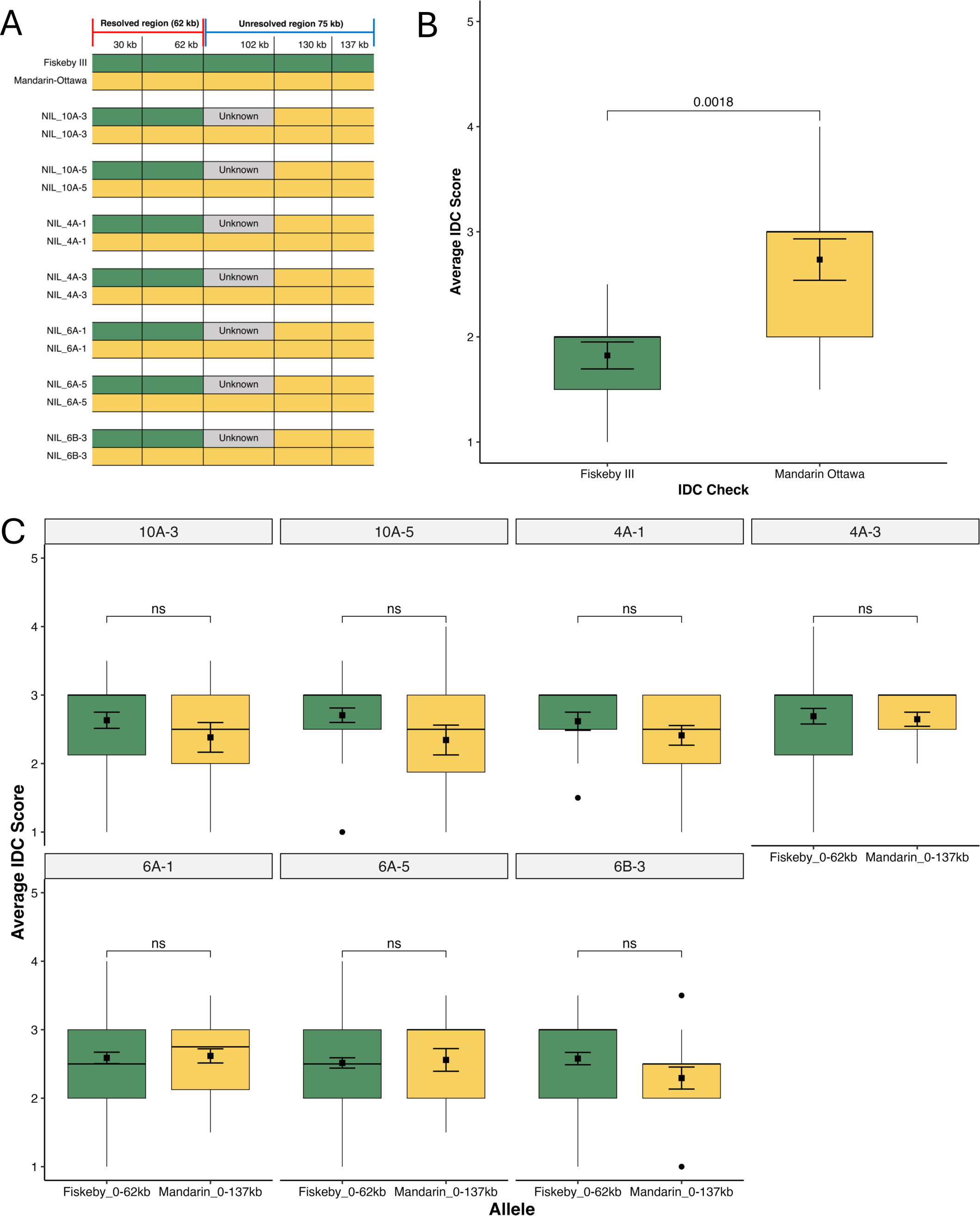
(A) Genotype data of different near-isogenic lines (NILs) with newly identified recombination points using KASP markers. The regions 0-62kb indicate allele differences carried by the NIL families.; (B) Parental lines used as IDC controls with Fiskeby III as resistant check and Mandarin (Ottawa) as susceptible check (C) A comparison of different alleles within each NIL family. No significant differences were observed between NILs carrying the Mandarin (Ottawa) allele from 0-137 kb and NILs carrying the Fiskeby III allele from 0-62 kb.

Figure 3C and Supplemental Figure 5 shows the result of the IDC screening of the seven different NIL families. Fiskeby III and Mandarin (Ottawa) were included as IDC checks, with Fiskeby III exhibiting a significantly lower IDC score than Mandarin (Ottawa), consistent with expectations (Figure 3B). Analysis of variance for the IDC scores (Supplemental Table 6) showed that environment, replication, allele, and replication within the environment were significant sources of variation. Importantly, field comparisons of the NIL pairs did not show significant differences between the lines with the Mandarin_0-137kb allele and the lines with the Fiskeby0_62kb allele (Figure 3C). All lines exhibited IDC susceptibility. It was observed that all seven families exhibited similar IDC responses in the field, cross-validating one another. Thus, it was concluded based on the two years of phenotypic testing that the IDC resistance gene is located within the interval Gm05: 62kb-137kb. This 75-kb interval contains 11 gene models based on the Wm82.a6 gene annotation.

### Comparative analysis of the Wm82.a6 and the ‘Fiskeby III’ genome assemblies within the fine-mapped IDC resistance region

We compared the two newly assembled genomes, Wm82.a6 and ‘Fiskeby III’, to detect sequence variation within the fine-mapped interval of the chromosome 05 IDC resistance QTL. Aligning the 75 kb interval between Wm82.a6 and Fiskeby III, single nucleotide polymorphisms, indels, and structural variants were detected. An approximately 10 kb insertion was found in the ‘Fiskeby III’ genome region (Figure 4A) within the aligned interval. The gene models were also visualized to identify the location of the insertions. The annotation of the inserted sequences identified two copies of a L1- 13 transposon (around 1,400 bp per copy) inserted within the gene model GlymaFiskIII.05G001200.1 and three copies of Gm-MULE34 transposon (around 2,300 bp per copy) located in between the two gene models GlymaFiskIII.05G001400.1 and GlymaFiskIII.05G001500.1 (Figure 4B). These two types of transposable elements were not observed within the fine-mapped region in the Wm82.a6 assembly.

**Figure 4.**
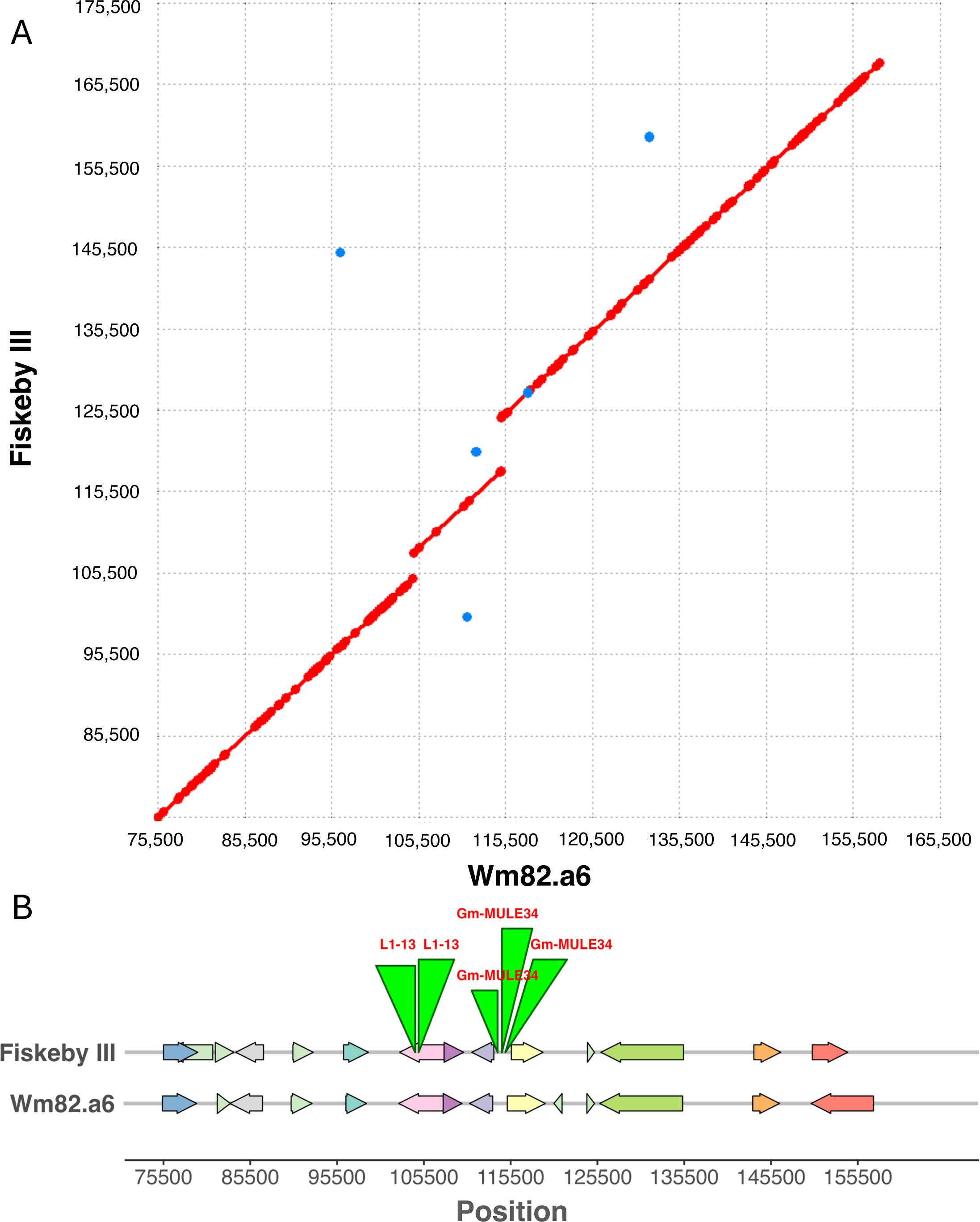
(A) Dot plots illustrating the maximal unique matches (MUMs) created by applying MUMmer 4.0 to the Fiskeby III and Wm82.a6 for the fine-mapped region based on the SNP marker at 62kb and 145kb. Red dots represent forward MUMs, while blue dots indicate reverse MUMs. (B) Gene models within the fine-mapped regions, annotated both in Fiskeby III and Wm82.a6, respectively. The green triangles indicate the presence of different transposable elements found in the Fiskeby III genome but absent in Wm82.a6, as identified by the RepeatMasker annotation.

It is not clear whether these insertions impact the QTL *per se*. Retrotransposons are known to be stress-activated in plants (Orozco-Arias et al., 2019) and the integration of transposable elements can also impact the expression of the adjacent genes. Furthermore, transposon integration may be accompanied by epigenetic changes (e.g., DNA methylation) of the insertion site (Orozco-Arias et al., 2019). These influences may lead to potential hypotheses of genomic impacts on the effect of the IDC QTL.

This is just one case study of how these newly assembled genomes can be used to develop molecularly mapped genetic markers, and lead to the development of functional genomics hypotheses. Numerous additional applications may be realized using these tools, particularly by exploiting the publicly available resequencing data resources of soybean (Liu et al., 2020; Torkamaneh et al., 2021; Valliyodan et al., 2021). Alignments of resequenced data to the gapless genomes and be utilized for allele mining (Chan et al., 2023), comparative genomics, diversity analysis, and genome wide association studies.

## CONCLUSIONS

In conclusion, we successfully assembled and annotated version 6 of Wm82 using a sub-line Wm82-ISU-01. The seeds that matched the Wm82.a6 were deposited in the soybean germplasm collection under PI 704477. This provides the soybean community with direct access to the sequenced line, making it the first such resource for comparative and functional genomics studies for the soybean research community. Our comparison of the three newer Wm82 assemblies unveiled previously unknown heterogeneity on chromosome 12, in addition to providing greater resolution to the previously identified introgressions on chromosomes 03 and 07. This is essential information for researchers investigating the different haplotypes in these genomic areas. Moreover, we utilized this genome for studying variations in the IDC resistance QTL locus, showcasing one of the many new utilities near-gapless genomes offer to the soybean community. In the era of gapless genome assembly, we assume that more high-quality assemblies will emerge in the near future. It is crucial for the soybean research community to have access to these shared resources. Establishing a shared gene annotation and nomenclature is essential, enabling the community to track genes from previous versions into the newer and more complete assemblies.

## MATERIALS AND METHODS

### Plant material, de novo assembly, and gene prediction

The soybean line Wm82-ISU-01 is a sub-line derived from a single plant of the soybean cultivar ‘Williams 82’ (Haun et al. 2011). The original seed source for Wm82-ISU-01 was Iowa State University (Haun et al. 2011). DNA from a single plant served as the source for the development of the Wm82.a6.v1 genome assembly. Leaf samples were collected, and high molecular weight DNA was extracted using the Qiagen HMW DNA Kit. The genome was assembled from 47x single haplotype Pacific BioSciences HiFi sequencing with an average read length of 17,049 bp. The assembly was performed and misjoins were identified using HI-C data. The JUICER pipeline was used to order, orient, and join contigs into chromosomes. The homozygous SNPs and indels for the final released sequence were corrected using 400 bp insert, 2x 150 Illumina reads with 57x coverage. For the gene annotation, RNA-seq was performed with RNA samples from different tissues of Williams 82 (Sreedasyam et al., 2023). Transcript assemblies were made from 2x 150 bp stranded paired-end Illumina RNA-seq reads using PERTRAN, which conducts genome guided transcriptome short read assembly via GSNAP (Wu and Nacu, 2010). Genes were predicted from transcriptome alignments, and using various homology-based predictors including EXONERATE (Slater and Birney, 2005), FGENESH+ (Salamov and Solovyev, 2000), FGENESH_ES, and PASA (Haas et al., 2003).

All the analyses of Wm82.a6 for visualizing gene and repeat positions were performed using GENESPACE (v 1.3.1) (Lovell et al., 2022) in R 4.3.1. The synteny plot was generated by aligning non-overlapping 1-kb windows reciprocally between the six Wm82 genomes with Wm82.a6 as reference using minimap2 with a kmer size of 25, window size of 20 and the “asm5” alignment preset, and parsing these two syntenic blocks of at least 40 windows. Telomeres, which are found on all 40 chromosomal termini, were inferred by mapping dense regions of telomere-specific kmers (CCCGAAA and CCCTAAA) and their reverse complements and clustering regions within 20 kb of chromosome termini that had at least 125 bases of exact matches to the kmers separated by no more than 100 bases of non-telomere sequence, which when combined had at least 80% telomere sequence. Cen 91/92 repeats (CentGm-2 and CentGm-1 respectively) were taken from (Gill et al., 2009) as CATTTGAATTTCTCGAGAGCTTCCGTTGTTCAATTTCGAGCGTCTCGATATATTATGCGCCTGAATCGGACCTCCGAGTTAAAAGTTATGAC and CGTTTGAATTTGCTCAGAGCTTCAGTATTCAATTTCGAGCGTCTCGATATATTACGGGACTCAATCAGACATCCGAGTAAAAAGTTATTGT. Instances of these repeats were identified in the genome assemblies by homology, using blastn with word-size = 11, soft masking off, ≥ 90% identity, and match length >= 85 bp. Hits overlapping by less than 10 bps were counted as distinct.

### Assessment of gene model characteristics and quality

BUSCO scores (Benchmarking Universal Single-Copy Orthologs; (Manni et al., 2021)) were calculated using BUSCO version 5.4.3, database fabales_odb10, hmmsearch 3.1. using protein sequences from the respective annotations (Wm82.a5 and Wm82.a6, Wm82_NJAU). Gene enrichment analyses were calculated using the GlycineMine tool at https://mines.legumeinfo.org/glycinemine/begin.do, with input gene lists consisting of genes present or absent in pairwise comparisons between the more complete Wm82 annotations, with Holm-Bonferroni correction for multiple testing. Gene correspondences were calculated for the 57 annotation sets in Glycine available at SoyBase as of February, 2024, using the Pandagma pangene workflow (https://github.com/legumeinfo/pandagma). The Wm82 gene correspondences (Supplemental Table 3) are a subset of the pangene set at https://data.legumeinfo.org/Glycine/GENUS/pangenes/Glycine.pan5.MKRS/.

### Comparative analysis of heterogeneity among the different Wm82 reference genomes

Previous research from (Haun et al., 2011) revealed genomic heterogeneity among different sub-lines of Wm82, primarily caused by differential introgressions of ‘Kingwa’ segments into the ‘Williams’ background during the ‘Williams 82’ breeding process. To test if the recent versions of the Wm82 genome assemblies also contained previously reported introgressions, we utilized the 50K SNP data from Wm82 parental lines Williams (PI548631) and Kingwa (PI548359). The genome positions of the 50K SNPs were identified for all the genome versions (Supplemental Table 7). Initially, the flanking sequences for the 50K SNPs from (Song et al., 2013) were aligned to the new reference genome using Minimap2 (Li, 2018) following the command “minimap2 -ax”.

The SNP state (matching ‘Williams’, matching ‘Kingwa’, or matching neither ‘Williams’ nor ‘Kingwa’) for each polymorphic 50K SNP was identified for each version of the Wm82 reference genome. The 50K SNP calls were used to visualize the ‘Kingwa’ introgression state for each genome version. The genome reference calls for each SNP position were compared with the data of ‘Williams’ and ‘Kingwa’ and categorized into four groups, the reference genome matching: (1) the ‘Williams’ SNP, (2) the ‘Kingwa’ SNP, (3) neither ‘Williams’ nor ‘Kingwa’, and (4) the reference genome is not polymorphic between the reference genome, ‘Williams’, and ‘Kingwa’.

Since the 50K SNP platform gives only a snapshot of the actual genomic heterogeneity among the Wm82 sub-lines, we examined the full set of SNP and indel sequence polymorphisms between the three new near gapless assemblies. Wm82.a6 was used as the reference then compared to Wm82.a5 (Garg et al., 2023) and Wm82.NJAU (Wang et al., 2023), respectively, as the query sequences. The SNPs calls from three genomes were then compared to determine which variant at a certain position is unique to a specific genome build. MUMmer4 (Marçais et al., 2018) was used for alignment following the parameters “--mum -l 40 -c 90” and filtered the SNPs and INDELs using “delta-filter -m -i 90 -l 100” and “show-snps -ClTr”. For all the SNPs variants, we calculated the SNP density in a 100 kb sliding window.

### Fine-mapping of iron deficiency chlorosis resistance in soybean

The materials for fine mapping were derived from the heterogeneous inbred families (HIFs) of a ‘Mandarin(Ottawa)’ x ‘Fiskeby III’ mapping population (Merry et al., 2019). The HIFs segregated for a 137 kb interval and were used to derive near isogenic lines (NILs) following (Tuinstra et al., 1997). Utilizing the ‘Fiskeby III’ genome assembly SNP data, five Kompetitive Allele Specific Primer (KASP) marker assays were designed around the region to screen positions within and around the segregating region (Gm05_30kb, Gm05_62kb, Gm05_102kb, Gm05_130kb, and Gm05_145kb) to identify new recombinants (Supplemental Table 8). After genotyping the HIFs, seven NIL families with recombination around the markers Gm05_62kb and Gm05_102kb were selected. Within each NIL family was a pair composed of two alleles, with one having the ‘Fiskeby III’ type allele from marker 0-62kb (Fiskeby_0-62kb) and the other having ‘Mandarin(Ottawa)’ type allele from 0-137 kb (Mandarin_0-137kb). After generating the seven NIL families, these lines were planted in fields prone to IDC, hereafter referred to as “IDC nurseries”. IDC nurseries were planted and managed at three Minnesota locations in 2022 (Danvers, Climax, and Foxhome), while two Minnesota locations were planted and managed in 2023 (Crookston and Climax). Plots consisted of single rows 91.4 cm in length spaced 76.2 cm apart. Plots were arranged in a matched pair randomized complete block design with three replications at each location. The entries that were paired and planted next to one another were the pairs of NILs within a family. This design was used in order to minimize spatial variability between the treatment comparisons of interest. The plots were scored for IDC symptoms using a 1-5 visual rating scale (Merry et al., 2022) two times per season with a two week interval between scoring dates. In addition to the NIL pairs, the parental lines were also grown in these trials, including ‘Mandarin(Ottawa)’ as susceptible and ‘Fiskeby III’ as resistant checks, respectively. The phenotypic data were analyzed using analysis of variance (ANOVA) to detect significant differences in IDC scores within each NIL family.

### Comparative analysis of ‘Wm82.a6’ assembly with the ‘Fiskeby III’ genome within the fine-mapped IDC resistance region

The genome was assembled for the IDC donor parent ‘Fiskeby III’ (PI 438471). Seeds for ‘Fiskeby III’ were obtained from the U.S. Department of Agriculture Germplasm Resource Information Network (GRIN). High molecular weight DNA extraction was performed using the Qiagen HMW DNA Kit. In brief, ‘Fiskeby III’ was assembled using 133.95x long-read PacBio coverage with an average read length of 11,253 bp. Using the Wm82.a4 assembly, misjoins were identified and polished. Scaffolds were oriented, ordered, and joined using Wm82.a4 synteny and Hi-C scaffolding. All the homozygous SNPs and indels in the released sequence were corrected using 50x Illumina reads. To predict gene models, RNA-seq was performed with RNA samples from different tissues of ‘Fiskeby III’ and sequenced using 2x150 Illumina sequencing. From around 2.6 billion paired reads, transcriptomes were assembled using PERTRAN (Wu and Nacu, 2010). Genes were predicted from transcriptome alignments, and using various homology-based predictors including EXONERATE(Slater and Birney, 2005), FGENESH+(Salamov and Solovyev, 2000), FGENESH_ES, and PASA, similar to the Wm82.a6 gene annotation.

The sequences for the narrowed 75 kb interval which spanned from the markers Gm05_62kb to Gm05_145kb were pulled out from the ‘Fiskeby III’ and Wm82.a6 genome assemblies found in Phytozome V13 (https://phytozome-next.jgi.doe.gov/info/Gmax_Wm82_a6_v1, https://phytozome-next.jgi.doe.gov/info/GmaxFiskeby_v1_1). The two genomes were aligned within the fine-mapped interval using MUMmer4 (Marçais et al., 2018) following the parameters “mummer -mum -b -c”, with Wm82.a6 as the reference sequence and ‘Fiskeby III’ as the query sequence. The resulting alignment was visualized using “mummerplot --png”.

Gene models within the aligned interval were determined from the Wm82.a6 annotation and repeats were identified using RepeatMasker annotation. The gene models and repeats were visualized with gggenomes (v0.9.9.9000) in R 4.3.1.

## ACKNOWLEDGMENTS AND FUNDING

The authors thank Adam Mahan and Allison Level for facilitating the rapid introduction of Wm82-ISU-01 (PI 704477) into the U.S. Department of Agriculture Germplasm Resource Information Network. The authors also thank Jacqueline Campbell and Rex Nelson for incorporating the Wm82.a6 assembly and annotation into the SoyBase database. The authors thank the University of Minnesota Doctoral Dissertation Fellowship program for supporting MJCE. The work (proposal: 10.46936/10.25585/60000843 and 10.46936/10.25585/60007936) conducted by the U.S. Department of Energy Joint Genome Institute (https://ror.org/04xm1d337), a DOE Office of Science User Facility, is supported by the Office of Science of the U.S. Department of Energy operated under Contract No. DE-AC02-05CH11231. Sequencing of the ‘Fiskeby III’ genome was supported by the United Soybean Board (project 1920- 172-0116-C to RMS and AJL). Fine-mapping of the IDC QTL was supported by the United Soybean Board (project 2311-209-0501 to RMS and AJL) and the United States Department of Agriculture (NIFA project 2022-67013-36128 to RMS and AJL). This work was also supported by the United States Department of Agriculture, Agricultural Research Service (USDA-ARS) CRIS Project 5030-21000-071-000D. The USDA is an equal opportunity provider and employer. Mention of trade names or commercial products in this article is solely for the purpose of providing specific information and does not imply recommendation or endorsement by the U.S. Department of Agriculture. This research used resources provided by the SCINet project and/or the AI Center of Excellence of the USDA Agricultural Research Service, ARS project numbers 0201- 88888-003-000D and 0201-88888-002-000D.

## AUTHOR CONTRIBUTIONS

MJCE, AJL, JS, and RMS designed the study. JJ, JW, LB, JT, and JG performed genome sequencing and chromosome-scale assemblies. SS, AS, TB, DG, and GS performed structural and functional annotation, and data release. MJCE, JTL, BDJ, SBC, JS, and RMS performed comparative genomic analyses. MJCE and AJL performed IDC fine-mapping analyses. All authors read and approved the final manuscript. MJCE, JTL, and RMS prepared the manuscript with input from all authors.

## SUPPLEMENTAL FIGURE LEGENDS

Supplemental Figure 1. The contig continuity map of the six versions of Wm82 genome builds. Each cycle through colors is 20 contigs containing 19 gaps. Contigs are defined as subsequences in the chromosomes separated by 10Ns in the older assemblies (a1 and a2) and 10,000Ns in the newer versions (a4, a5, NJAU, and a6). The telomere density was set at 80%.

Supplemental Figure 2. Analysis of Kingwa introgressions in six different Williams 82 reference genome builds based on Infinium 50K SNP genotypes. The SNP positions were mapped against the six versions of reference genomes. Blue dots indicate SNPs matching Williams, red dots indicate SNPs matching Kingwa, green dots indicate SNPs matching neither Williams nor Kingwa, and gray ‘X’ represents non-polymorphic SNPs between Williams and Kingwa.

Supplemental Figure 3. SNP density per 100 kb window for SNPs unique to Wm82.a5 and SNPs unique to Wm82.NJAU compared to Wm82.a6 across 20 chromosomes.

Supplemental Figure 4. Fiskeby III contig continuity plot. Each cycle through colors is 20 contigs containing 19 gaps. Contigs are defined as subsequences in the chromosomes separated by 10,000 Ns and telomere density was set at 80%.

Supplemental Figure 5. Comparison of means for the new recombinants across the seven families.

## SUPPLEMENTAL TABLE LEGENDS

Supplemental Table 1. Centromeric repeat abundance for each chromosome in the Wm82 genome assemblies and the Fiskeby III genome assembly. Values represent the number of repeat units on each chromosome based on the canonical 91/92-bp soybean centromeric repeat units as described in Gill et al. (2009).

Supplemental Table 2. Telomeric repeat abundance at the chromosome start or end for each Wm82 genome assembly and the Fiskeby III genome assembly. Values represent the number of repeat units on each chromosome based on the canonical 7-bp plant telomeric repeat unit.

Supplemental Table 3. Gene predictions for the six Wm82 genome assembly versions. Each row is based on the assigned pangene available at SoyBase as of February, 2024. “NONE” indicates that the annotation for the given assembly version did not predict a gene that matches this pangene.

Supplemental Table 4. Number of indels in detected in Wm82.a5 and Wm82.NJAU compared to Wm82.a6.

Supplemental Table 5. Genome assembly metrics of Fiskeby III.

Supplemental Table 6. Analysis of variance for IDC scores of the near isogenic lines for fine mapping.

Supplemental Table 7. Liftover positions of the SoySNP50K for six different Wm82 reference genome assemblies. “NA” indicates that a specific SNP did not map in a given genome assembly.

Supplemental Table 8. KASP markers used for genotyping IDC fine-mapping recombinants.

## REFERENCES

Belser C, Baurens FC, Noel B, Martin G, Cruaud C, Istace B, Yahiaoui N, Labadie K, Hřibová E, Doležel J, et al. Telomere-to-telomere gapless chromosomes of banana using nanopore sequencing. Commun Biol. 2021:4(1):1047. doi: 10.1038/s42003-021-02559-3

Bernard RL, Cremeens C. Registration of “Williams 82” soybean. Crop Sci. 1988:28(6):1027–1028. doi: 10.2135/cropsci1988.0011183X002800060049x

Burton AL, Burkey KO, Carter TE Jr, Orf J, Cregan PB. Phenotypic variation and identification of quantitative trait loci for ozone tolerance in a Fiskeby III × Mandarin (Ottawa) soybean population. Theor Appl Genet. 2016:129(6):1113–1125. doi: 10.1007/s00122-016-2687-1

Butenhoff K. QTL mapping and GWAS identify sources of iron deficiency chlorosis and canopy wilt tolerance in the Fiskeby III x Mandarin (Ottawa) soybean population. 2015:Master of Science. University of Minnesota, Saint Paul, MN, United States. https://hdl.handle.net/11299/170730

Chan YO, Dietz N, Zeng S, Wang J, Flint-Garcia S, Salazar-Vidal MN, Škrabišová M, Bilyeu K, Joshi T. The Allele Catalog Tool: a web-based interactive tool for allele discovery and analysis. BMC Genomics. 2023:24(1):107. doi: 10.1186/s12864-023-09161-3

Chen J, Wang Z, Tan K, Huang W, Shi J, Li T, Hu J, Wang K, Wang C, Xin B, et al. A complete telomere-to-telomere assembly of the maize genome. Nat Genet. 2023:55(7):1221-1231. doi: 10.1038/s41588-023-01419-6

Deng Y, Liu S, Zhang Y, Tan J, Li X, Chu X, Xu B, Tian Y, Sun Y, Li B, et al. A telomere-to-telomere gap-free reference genome of watermelon and its mutation library provide important resources for gene discovery and breeding. Mol Plant. 2022:15(8):1268–1284. doi: 10.1016/j.molp.2022.06.010

Do TD, Vuong TD, Dunn D, Smothers S, Patil G, Yungbluth DC, Chen P, Scaboo A, Xu D, Carter TE, et al. Mapping and confirmation of loci for salt tolerance in a novel soybean germplasm, Fiskeby III. Theor Appl Genet. 2018:131(3):513–524. doi: 10.1007/s00122-017-3015-0

Fasoula VA, Boerma HR (2007) Intra-cultivar variation for seed weight and other agronomic traits within three elite soybean cultivars. Crop Sci. 2007:47(1): 367–373. doi: 10.2135/cropsci2005.09.0334

Garg V, Khan AW, Fengler K, Llaca V, Yuan Y, Vuong TD, Harris C, Chan TF, Lam HM, Varshney RK, et al. Near-gapless genome assemblies of Williams 82 and Lee cultivars for accelerating global soybean research. Plant Genome. 2023:16(4):e20382. doi: 10.1002/tpg2.20382

Gill N, Findley S, Walling JG, Hans C, Ma J, Doyle J, Stacey G, Jackson SA. Molecular and chromosomal evidence for allopolyploidy in soybean. Plant Physiol. 2009:151(3):1167–1174. doi: 10.1104/pp.109.137935

Gladman N, Goodwin S, Chougule K, McCombie RW, Ware D. Era of gapless plant genomes: innovations in sequencing and mapping technologies revolutionize genomics and breeding. Curr Opin Biotechnol. 2023:79:102886. doi: 10.1016/j.copbio.2022.102886

Haas BJ, Delcher AL, Mount SM, Wortman JR, Smith RK Jr, Hannick LI, Maiti R, Ronning CM, Rusch DB, Town CD, et al. Improving the Arabidopsis genome annotation using maximal transcript alignment assemblies. Nucleic Acids Res. 2003:31(19):5654–5666. doi: 10.1093/nar/gkg770

Haun WJ, Hyten DL, Xu WW, Gerhardt DJ, Albert TJ, Richmond T, Jeddeloh JA, Jia G, Springer NM, Vance CP, et al. The composition and origins of genomic variation among individuals of the soybean reference cultivar Williams 82. Plant Physiol. 2011:155(2):645–655. doi: 10.1104/pp.110.166736

Huang X. A complete telomere-to-telomere assembly provides new reference genome for rice. Mol Plant. 2023:16(9):1370–1372. doi: 10.1016/j.molp.2023.08.007

Li H. Minimap2: pairwise alignment for nucleotide sequences. Bioinformatics. 2018:34(18):3094–3100. doi: 10.1093/bioinformatics/bty191

Liu Y, Du H, Li P, Shen Y, Peng H, Liu S, Zhou GA, Zhang H, Liu Z, Shi M, et al. Pan-Genome of wild and cultivated soybeans. Cell. 2020:182(1):162–176.e13. doi: 10.1016/j.cell.2020.05.023

Lovell JT, Sreedasyam A, Schranz ME, Wilson M, Carlson JW, Harkess A, Emms D, Goodstein DM, Schmutz J. GENESPACE tracks regions of interest and gene copy number variation across multiple genomes. Elife. 2022:11:e78526. doi: 10.7554/eLife.78526

Manni M, Berkeley MR, Seppey M, Zdobnov EM. BUSCO: Assessing genomic data quality and beyond. Curr Protoc. 2021:1(12):e323. doi: 10.1002/cpz1.323

Marçais G, Delcher AL, Phillippy AM, Coston R, Salzberg SL, Zimin A. MUMmer4: A fast and versatile genome alignment system. PLoS Comput Biol. 2018:14(1):e1005944. doi: 10.1371/journal.pcbi.1005944

Merry RA. Fine-mapping, physiological evaluation, and candidate gene exploration of an iron deficiency chlorosis tolerance locus in soybean. 2020:University of Minnesota, Saint Paul, MN, United States. https://hdl.handle.net/11299/216359

Merry R, Butenhoff K, Campbell BW, Michno JM, Wang D, Orf JH, Lorenz AJ, Stupar RM. Identification and fine-mapping of a soybean quantitative trait locus on chromosome 5 conferring tolerance to iron deficiency chlorosis. Plant Genome. 2019:12(3):1–13. doi: 10.3835/plantgenome2019.01.0007

Merry R, Dobbels AA, Sadok W, Naeve S, Stupar RM, Lorenz AJ. Iron deficiency in soybean. Crop Sci. 2022:62(1):36–52. doi: 10.1002/csc2.20661

Mihelich NT, Mulkey SE, Stec AO, Stupar RM. Characterization of genetic heterogeneity within accessions in the USDA soybean germplasm collection. Plant Genome. 2020:13(1):e20000. doi: 10.1002/tpg2.20000

Navrátilová P, Toegelová H, Tulpová Z, Kuo YT, Stein N, Doležel J, Houben A, Šimková H, Mascher M. Prospects of telomere-to-telomere assembly in barley: Analysis of sequence gaps in the MorexV3 reference genome. Plant Biotechnol J. 2022:20(7):1373–1386. doi: 10.1111/pbi.13816

Orozco-Arias S, Isaza G, Guyot R. Retrotransposons in plant genomes: Structure, identification, and classification through bioinformatics and machine learning. Int J Mol Sci. 2019:20(15):3837. doi: 10.3390/ijms20153837

Salamov AA, Solovyev VV. Ab initio gene finding in Drosophila genomic DNA. Genome Res. 2000:10(4):516–522. doi: 10.1101/gr.10.4.516

Schmutz J, Cannon SB, Schlueter J, Ma J, Mitros T, Nelson W, Hyten DL, Song Q, Thelen JJ, Cheng J, et al. Genome sequence of the palaeopolyploid soybean. Nature. 2010:463(7278):178-183. doi: 10.1038/nature08670

Sebastian SA, Streit LG, Stephens PA, Thompson JA, Hedges BR, Fabrizius MA, Soper JF, Schmidt DH, Kallem RL, Hinds MA, et al. Context-specific marker-assisted selection for improved grain yield in elite soybean populations. Crop Sci. 2010:50(4):1196–1206. doi: 10.2135/cropsci2009.02.0078

Slater GS, Birney E. Automated generation of heuristics for biological sequence comparison. BMC Bioinformatics. 2005:6:31. doi: 10.1186/1471-2105-6-31

Song Q, Hyten DL, Jia G, Quigley CV, Fickus EW, Nelson RL, Cregan PB. Development and evaluation of SoySNP50K, a high-density genotyping array for soybean. PLoS One. 2013:8(1):e54985. doi: 10.1371/journal.pone.0054985

Song Q, Hyten DL, Jia G, Quigley CV, Fickus EW, Nelson RL, Cregan PB. Fingerprinting soybean germplasm and its utility in genomic research. G3 (Bethesda). 2015:5(10):1999-2006. doi: 10.1534/g3.115.019000

Song Q, Jenkins J, Jia G, Hyten DL, Pantalone V, Jackson SA, Schmutz J, Cregan PB. Construction of high resolution genetic linkage maps to improve the soybean genome sequence assembly Glyma1.01. BMC Genomics. 2016:17:33. doi: 10.1186/s12864-015-2344-0

Sreedasyam A, Plott C, Hossain MS, Lovell JT, Grimwood J, Jenkins JW, Daum C, Barry K, Carlson J, Shu S, et al. JGI Plant Gene Atlas: an updateable transcriptome resource to improve functional gene descriptions across the plant kingdom. Nucleic Acids Res. 2023:51(16):8383–8401. doi: 10.1093/nar/gkad616

Torkamaneh D, Lemay MA, Belzile F. The pan-genome of the cultivated soybean (PanSoy) reveals an extraordinarily conserved gene content. Plant Biotechnol J. 2021:19(9):1852–1862. doi: 10.1111/pbi.13600

Tuinstra MR, Ejeta G, Goldsbrough PB. Heterogeneous inbred family (HIF) analysis: a method for developing near-isogenic lines that differ at quantitative trait loci. Theor Appl Genet 1997:95:1005–1011. doi: 10.1007/s001220050654

Valliyodan B, Brown AV, Wang J, Patil G, Liu Y, Otyama PI, Nelson RT, Vuong T, Song Q, Musket TA, et al. Genetic variation among 481 diverse soybean accessions, inferred from genomic re-sequencing. Sci Data. 2021:8(1):50. doi: 10.1038/s41597-021-00834-w

Valliyodan B, Cannon SB, Bayer PE, Shu S, Brown AV, Ren L, Jenkins J, Chung CY, Chan TF, Daum CG, et al. Construction and comparison of three reference-quality genome assemblies for soybean. Plant J. 2019:100(5):1066–1082. doi: 10.1111/tpj.14500

Varshney RK, Sinha P, Singh VK, Kumar A, Zhang Q, Bennetzen JL. 5Gs for crop genetic improvement. Curr Opin Plant Biol. 2020:56:190–196. doi: 10.1016/j.pbi.2019.12.004

Wang B, Yang X, Jia Y, Xu Y, Jia P, Dang N, Wang S, Xu T, Zhao X, Gao S, et al. High-quality Arabidopsis thaliana genome assembly with Nanopore and HiFi long reads. Genomics Proteomics Bioinformatics. 2022:20(1):4–13. doi: 10.1016/j.gpb.2021.08.003

Wang L, Zhang M, Li M, Jiang X, Jiao W, Song Q. A telomere-to-telomere gap-free assembly of soybean genome. Mol Plant. 2023:16(11):1711–1714. doi: 10.1016/j.molp.2023.08.012

Wu TD, Nacu S. Fast and SNP-tolerant detection of complex variants and splicing in short reads. Bioinformatics. 2010:26(7):873–881. doi: 10.1093/bioinformatics/btq057

Zhou Y, Zhang J, Xiong X, Cheng Z-M, Chen F. De novo assembly of plant complete genomes. Tropical Plants. 2022:1:7. doi: 10.48130/TP-2022-0007

