## Supplemental figures for "Assembly, comparative analysis, and utilization of a single haplotype reference genome for soybean"

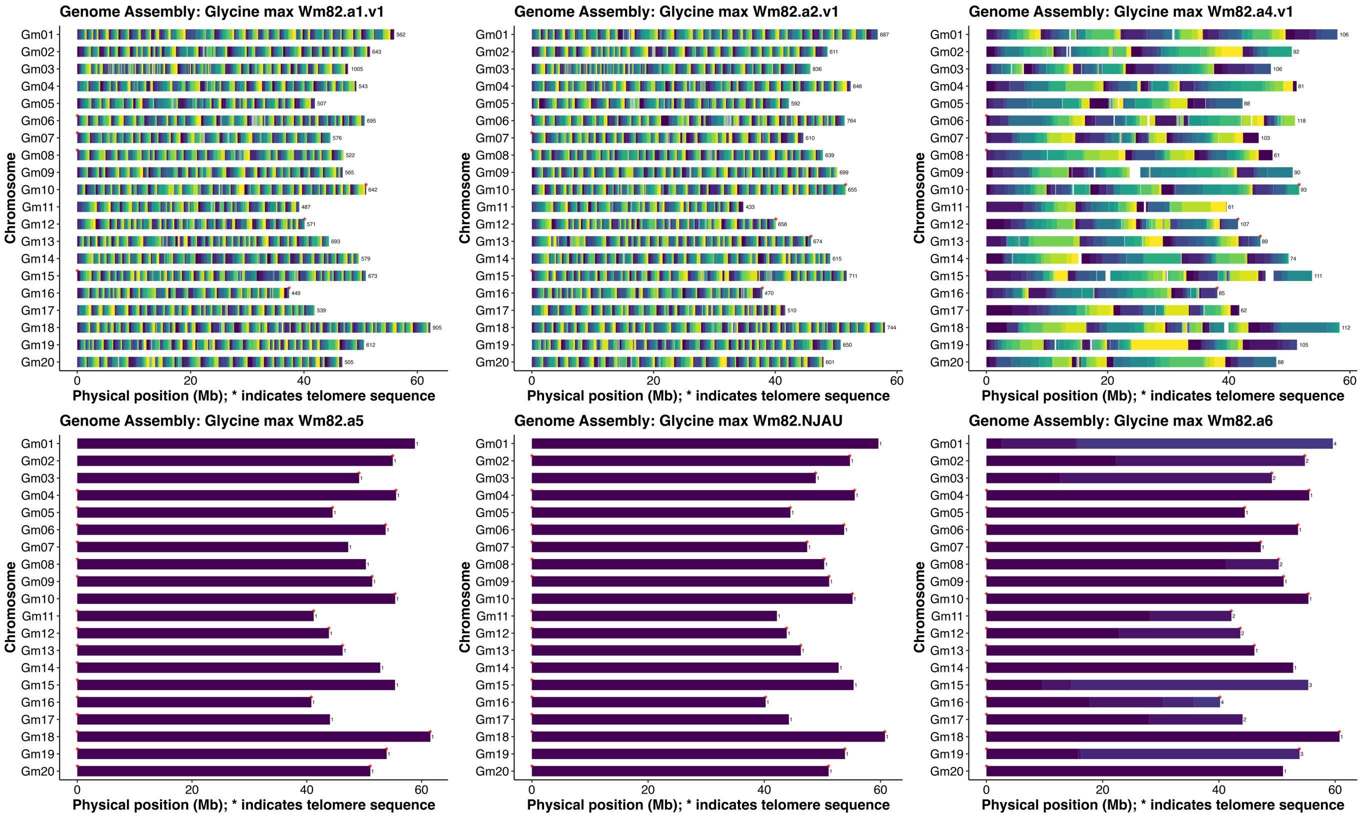


Supplemental Figure 1. The contig continuity map of the six versions of Wm82 genome builds. Each cycle through colors is 20 contigs containing 19 gaps. Contigs are defined as subsequences in the chromosomes separated by 10Ns in the older assemblies (a1 and a2) and 10,000Ns in the newer versions (a4, a5, NJAU, and a6). The telomere density was set at 80%.

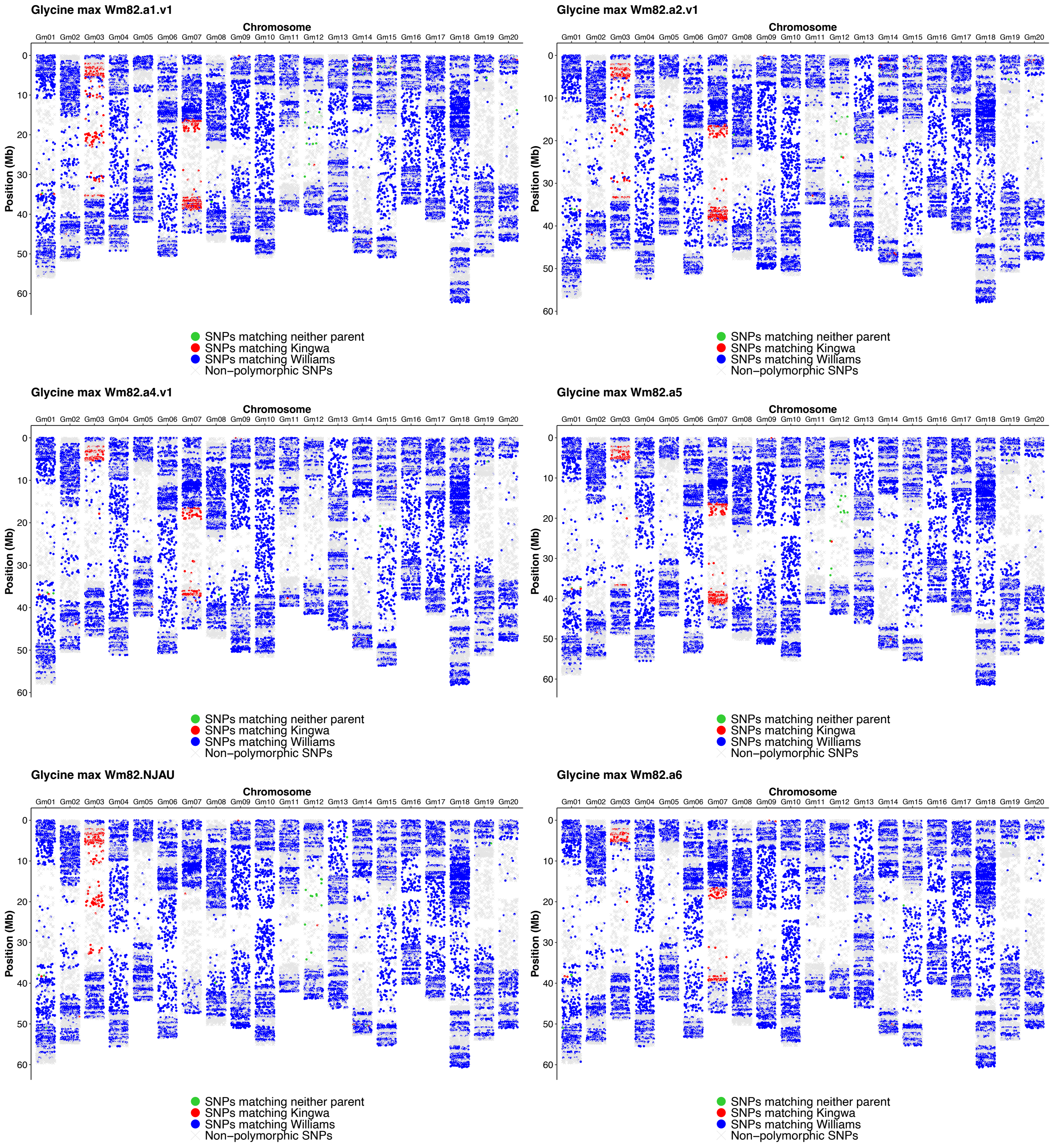


Supplemental Figure 2. Analysis of Kingwa introgressions in six different Williams 82 reference genome builds based on Infinium 50K SNP genotypes. The SNP positions were mapped against the six versions of reference genomes. Blue dots indicate SNPs matching Williams, red dots indicate SNPs matching Kingwa, green dots indicate SNPs matching neither Williams nor Kingwa, and gray ‘X’ represents non-polymorphic SNPs between Williams and Kingwa.

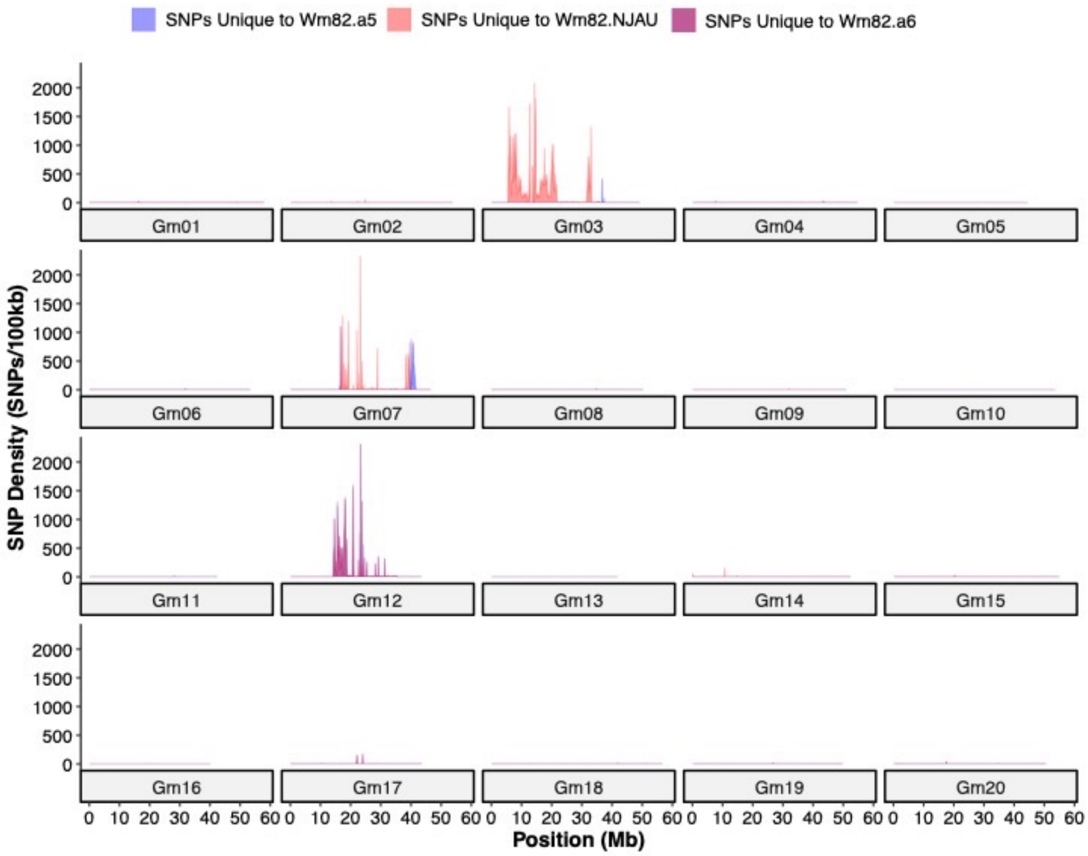


Supplemental Figure 3. SNP density per 100 kb window for SNPs unique to Wm82.a5 and SNPs unique to Wm82.NJAU compared to Wm82.a6 across 20 chromosomes.

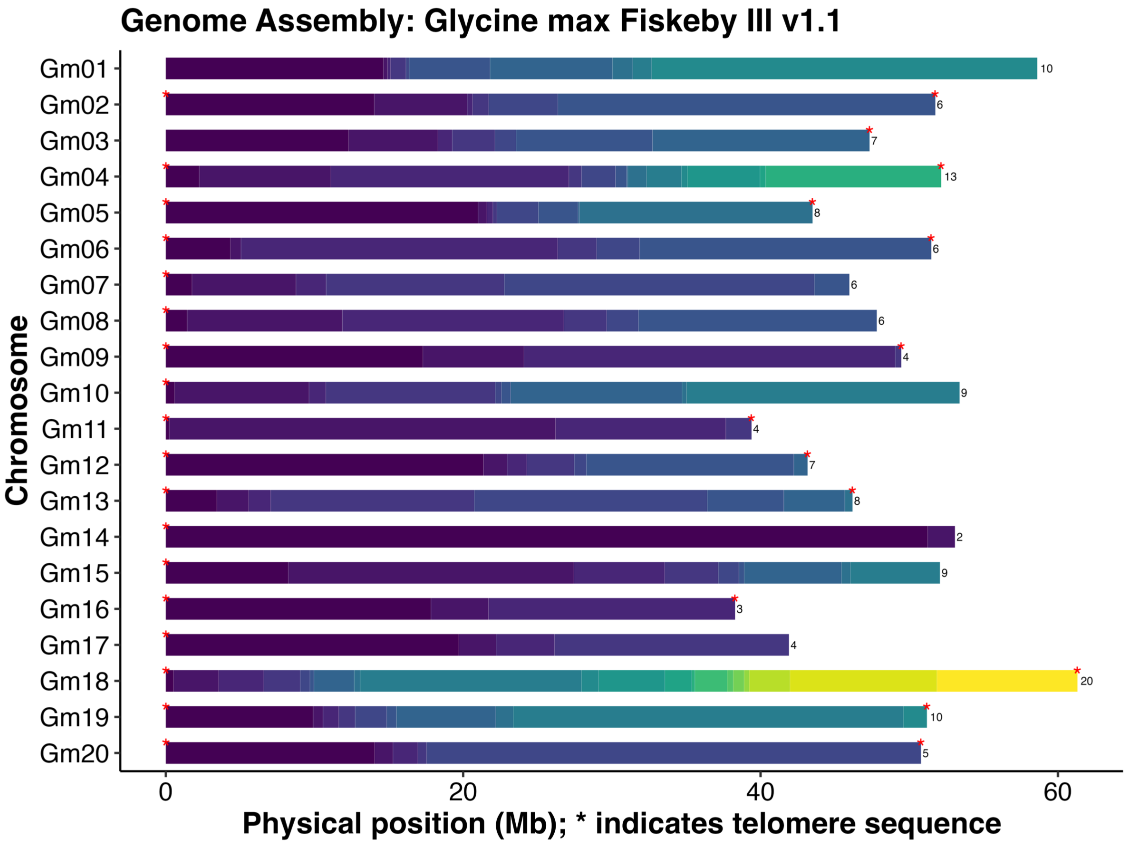


Supplemental Figure 4. Fiskeby III contig continuity plot. Each cycle through colors is 20 contigs containing 19 gaps. Contigs are defined as subsequences in the chromosomes separated by 10,000 Ns and telomere density was set at 80%.

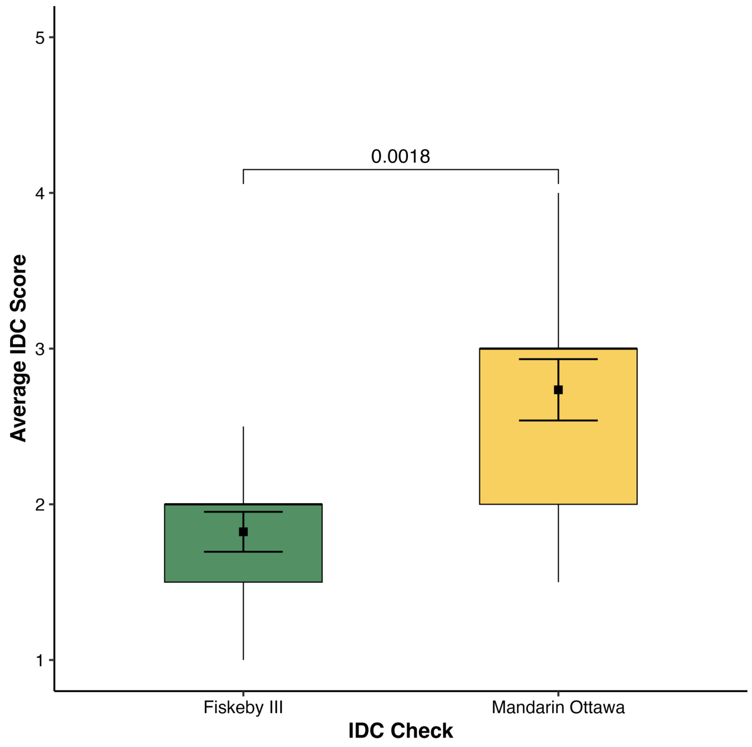


Supplemental Figure 5. Comparison of means for the new recombinants across the seven families.
